# Hypoxia and loss of *GCM1* expression prevents differentiation and contact inhibition in human trophoblast stem cells

**DOI:** 10.1101/2024.09.10.612343

**Authors:** Jessica K. Cinkornpumin, Sin Young Kwon, Anna-Maria Prandstetter, Theresa Maxian, Jacinthe Sirois, James Goldberg, Joy Zhang, Deepak Saini, Purbasa Dasgupta, Mariyan J. Jeyarajah, Stephen J. Renaud, Soumen Paul, Sandra Haider, William A. Pastor

## Abstract

The placenta develops alongside the embryo and nurtures fetal development to term. During the first stages of embryonic development, due to low blood circulation, the blood and ambient oxygen supply is very low (∼1-2% O_2_) and gradually increases upon placental invasion. While a hypoxic environment is associated with stem cell self-renewal and proliferation, persistent hypoxia may have severe effects on differentiating cells and could be the underlying cause of placental disorders. We find that human trophoblast stem cells (hTSC) thrive in low oxygen, whereas differentiation of hTSC to trophoblast to syncytiotrophoblast (STB) and extravillous trophoblast (EVT) is negatively affected by hypoxic conditions. The pro-differentiation factor GCM1 (human Glial Cell Missing-1) is downregulated in low oxygen, and concordantly there is substantial reduction of GCM1-regulated genes in hypoxic conditions. Knockout of GCM1 in hTSC caused impaired EVT and STB formation and function, reduced expression of differentiation-responsive genes, and resulted in maintenance of self-renewal genes. Treatment with a PI3K inhibitor reported to reduce GCM1 protein levels likewise counteracts spontaneous or directed differentiation. Additionally, chromatin immunoprecipitation of GCM1 showed enrichment of GCM1-specific binding near key transcription factors upregulated upon differentiation including the contact inhibition factor *CDKN1C.* Loss of *GCM1* resulted in downregulation of *CDKN1C* and corresponding loss of contact inhibition, implicating GCM1 in regulation of this critical process.

## Introduction

Placentation establishes the maternal-fetal interface that facilitates the successful progression of the pregnancy^1^. During the first major specification event at the early blastocyst stage, the embryo undergoes cell division and cavitation to form an outer layer of cells, termed the trophectoderm (TE) as well as the inner cell mass, which contains the epiblast that gives rise to the embryo proper^2^. Cells from the TE, upon implantation, give rise to cells called cytotrophoblasts (CTB) which can differentiate into the extravillous trophoblast (EVT) and syncytiotrophoblast (STB). These cells are organized into structures called villi, in which CTBs line the inside of the villus and STBs line the outside, mediating the exchange of nutrients, oxygen (O_2_), and waste. At the tips of the villi, the points of contact with maternal tissue, the CTBs form a cell column and differentiate into EVTs. Distinct subtypes of mature EVTs act to invade maternal decidua and to remodel spiral arteries, enabling proper blood flow to the placenta. Establishing this network of organized cell types and tissue structures is essential to maintain the connection between the mother and fetus, allowing for proper fetal health and development^3–8^.

During the first trimester of human pregnancy, some EVTs establish plugs blocking the uterine spiral arteries. For the initial stages of the first trimester, the conceptus develops in a low oxygen environment (∼2.5% O_2_, Hypoxia)^9,10^. After approximately eight weeks, the trophoblast plugs disintegrate and endovascular extravillous cytotrophoblasts (eEVT) invade the uterine spiral arteries where they degrade resident smooth muscle and endothelial cells (ECs) and replace them with a trophoblast cells.^11^ This expands the arterial lumen, provides blood to the placenta, and raises oxygen tension (∼8.6% O_2_)^9,12,13^.

Oxygen tension is clearly important in regulation of the trophoblast but how it regulates placental cell self-renewal and differentiation is not entirely clear. Aspects of response to hypoxia are universal. At high oxygen levels one of several prolyl hydroxylases will oxidize the HIF-family transcription factors, HIF1α and HIF2α. The oxidized prolines are then recognized by VHL which ubiquitinates the HIFs and targets them for destruction^14^. Low oxygen reduces the activity of the prolyl hydroxylases and thus stabilizes HIF-family transcription factors. HIF1α or HIF2α then dimerizes with the transcription factor ARNT and promote transcription of target genes. Certain HIF-targets are consistent across cell types, and hypoxia frequently promotes angiogenesis and a shift from oxidative respiration to glycolysis^14^. With regard to placenta, mice carrying deletions of *Hif1α/Hif2α* or *Arnt* both undergo midgestational embryonic lethality^15,16^. These mutants show reduced labyrinth vascularization, consistent with the classic role for hypoxia signaling in angiogenesis^15,16^. Intriguingly they also fail to maintain their spongiotrophoblast population (the murine functional equivalent to EVT precursors) and *Hif1a^−/−^Hif2a^−/−^* or *Arnt^−/−^* murine trophoblast stem cells (mTSCs) preferentially differentiate to STB rather than spongiotrophoblast lineage^15^. Loss of the prolyl hydroxylase *Phd2*, which causes elevated HIF-stability, results in reduced expression of STB markers and an increase in spongiotrophoblast^17^, while a *Vhl^−/−^*mouse features a loss of STB altogether^18^. Thus, a consistent feature in murine models is that hypoxia is unfavorable to STB differentiation but conducive for spongiotrophoblast.

In humans, hypoxia is well-established to block differentiation to STBs^19–22^. The effects on EVT differentiation are less clear. When explants of human placental villi are cultured in high oxygen, expression of ITGA1 (a mature EVT marker) is observed at the edges of the explant where cell column CTBs are found. In low oxygen, instead of ITGA1 expression, cell column CTBs proliferated and appeared to show elevated HLA-G^12^. Low O_2_ is reported to reduce CTB invasiveness and block expression of ITGA1, further supporting a role for oxygen in positive regulation of EVT differentiation^23^. Another study reports higher HLA-G upon culture of human CTB at low O_2_ and indicates a positive role for hypoxia in promoting conversion of CTB to less mature, proximal column-EVT^22^. With the discovery of culture conditions that allow for indefinite culture of CTBs *in vitro* as human trophoblast stem cells (hTSCs)^24^, we sought to determine the molecular and phenotypic effects of oxygen concentration on human placental cells.

## Results

### Hypoxia Maintains Trophoblast Stemness

To determine the effect of hypoxia on hTSC growth, we took hTSCs, which are commonly maintained in 20% O_2_, and cultured them in 20%, 5% and 2% O_2_. After the first 72 hours of culture, we performed flow cytometry for the hTSC cell surface markers ITGA6 and EpCAM, and the EVT markers ITGA1 and HLA-G (Fig. 1A). We observed noticeable depletion of ITGA1 and slightly increased HLA-G expression in the lower oxygen concentrations in several hTSC lines (Fig. 1A, 1B), similar to what was observed in explants by Genbacev and colleagues. Continued culture of these cells in their respective oxygen conditions resulted in near complete loss of ITGA1^hi^ expression cells in both 5% and 2% O_2_ (Fig. S1A). Similarly, hTSCs cultured in 20% O_2_ showed some spontaneous expression of the STB marker hCGB, which was reduced in low oxygen (Fig. S1B, S1C). Regions of dense cell-to-cell contact showed expression of the differentiation marker NOTCH1 in hTSCs at 20% O_2_, but not in lower oxygen (Fig. S1D). In addition to lower expression of differentiation markers, we observed higher cell density in lower oxygen culture conditions (Fig. S1B, S1E). Collectively, these results indicated that low oxygen aids in stemness and proliferation, while high oxygen promotes spontaneous differentiation.

**Figure 1.**
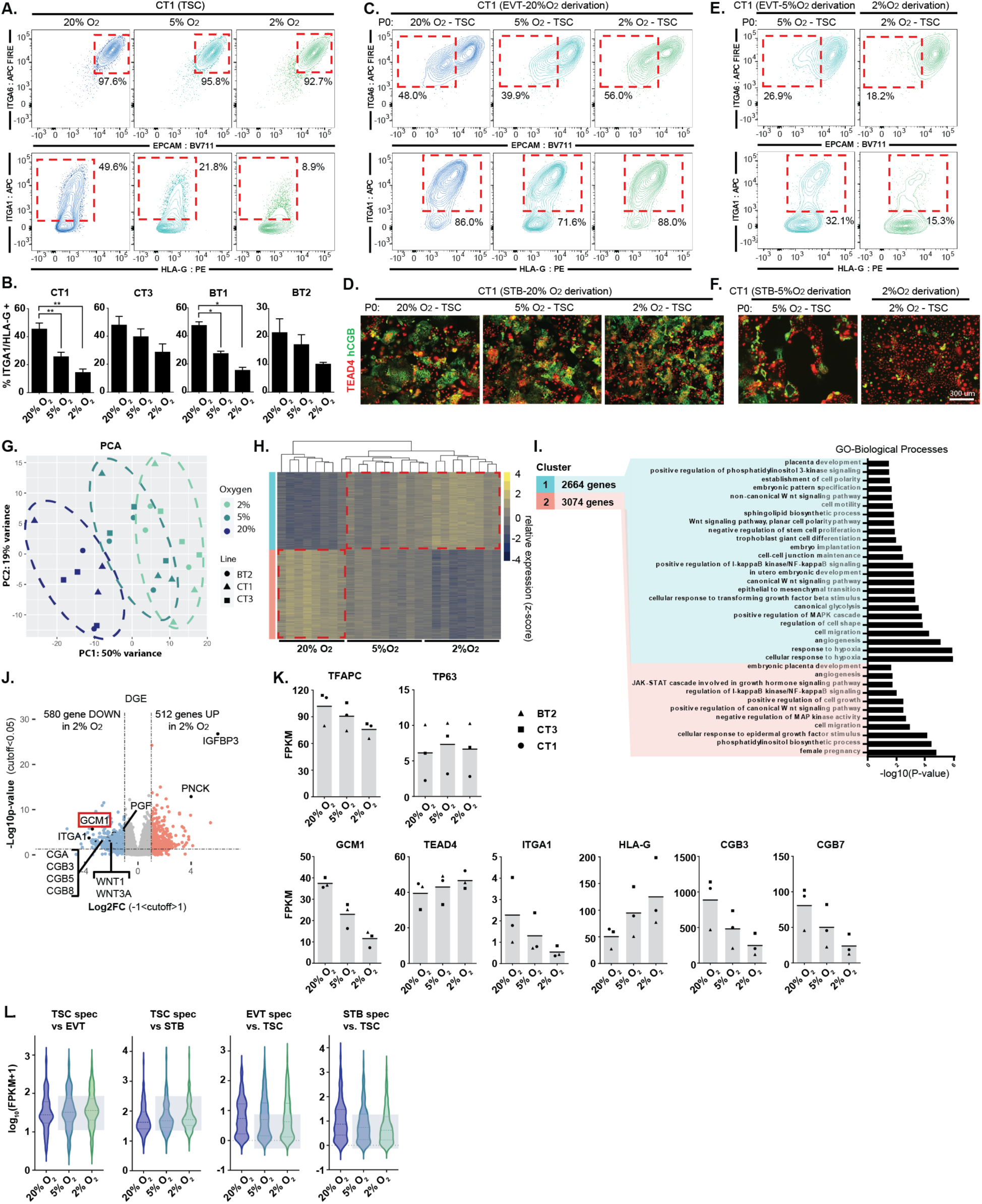
Reduced and impaired hTSC differentiation in hypoxic conditions. **A.** Trophoblast stem cells were cultured for 72hrs in varying levels of oxygen (20%, 5%, 2% O_2_). Flow cytometry plots indicate levels of hTSC (ITGA6, EPCAM) and EVT (ITGA1, HLA-G) markers. Note reduction in ITGA1^+^ population in low O_2_. **B.** ITGA1^+^ HLA-G^+^ population in O_2_ and cell line indicated (n=3 replicates for each cell line). **C.** EVT differentiation in 20% O_2_ starting with hTSC in oxygen concentration indicated. Successful differentiation is indicated by the upregulation of surface markers ITGA1 and HLA-G and downregulation of EPCAM and ITGA6. **D.** STB differentiation in 20% O_2_ starting with hTSC in oxygen concentration indicated. STB formation is indicated by loss of TEAD4 and increase in hCGB staining in a cell. **E.** EVT differentiation undertaken at oxygen level indicated. **F.** STB differentiation undertaken at oxygen level indicated. **G.** Principle component analysis (PCA) showing gene expression from hTSC cultured in varying oxygen concentrations. (3 cell lines; BT2, CT1, CT3; n=3 replicates for each line in each condition, except BT2 at 20% O_2_ n=2). **H.** Hierarchical gene clustering of RNA-seq samples in (G). Red dotted lines indication the shift in gene expression from 20% O_2_ and 2% O_2_ labeled as Cluster 1 and Cluster 2. **I.** GSEA analysis of Cluster 1 and Cluster 2. **J.** Volcano plot showing gene expression differences between TSCs cultured in 20% O_2_ to TSCs cultured in 2% O_2_. Dashed lines indicate significance and Log2 fold-change cutoff. **K.** Bar graphs showing FPKM of specific genes of interest (same samples as in G). **L.** Violin plot showing expression of genes specific to hTSC, EVT or STB for hTSCs grown in the indicated oxygen concentration.

Based on these findings, we cultured hTSCs and conducted directed differentiation at varying oxygen conditions. hTSC cultured in 20%, 5% or 2% O_2_ successfully differentiated to EVT or STB if differentiation was undertaken at 20% O_2_ (Fig. 1C, 1D). However, when we perform EVT or STB differentiation at reduced oxygen levels, we observed dramatic impairment of differentiation (Fig. 1E, 1F, S1F). EVTs differentiated in reduced oxygen failed to downregulate EpCAM or upregulate ITGA1 and HLA-G (Fig. 1E), while STBs in 2% O_2_ showed a higher percentage of cells retaining TEAD4 and a lower percentage expressing hCGB (Fig. 1F, S1F). Thus, oxygen promotes both spontaneous and directed differentiation of hTSCs.

### Trophoblast Differentiation Transcription Factor, GCM1, is Oxygen Sensitive

We performed RNA-sequencing (RNA-seq) on three hTSC lines (CT1, CT3, BT2; female) cultured in 20%, 5%, or 2% O_2_. Principle component analysis (PCA) and a correlation matrix show a strong dependence of gene expression on oxygen tension (Fig. 1G, S1G). Gene cluster and enrichment analysis^25^ indicate that the genes in both Cluster 1 (genes upregulated in 2% O_2_) and Cluster 2 (upregulated in 20% O_2_) showed general enrichment for various placental-related terms while only Cluster 1 corresponded to hypoxia response and WNT activation (Fig. 1H, 1I).

Genes associated with EVT and STBs expression such as *ITGA1*, *OVOL1* and various chorionic gonadotropin genes were among the genes downregulated in the 2% O_2_ condition (Fig. 1J, 1K, S1H, Table S1). Using published gene expression data, we identified 100 genes specific to EVT and STB differentiation and 100 genes specific to hTSC relative to these cell types^24^. hTSC genes were higher in 2% O_2_, while EVT and STB genes were lower, further indicating that hypoxia broadly suppresses genes associated with differentiation (Fig. 1L, Table S2). Interestingly, consistent with flow cytometry data (Fig. 1A), HLA-G was positively regulated by hypoxia (Fig. 1K), indicating that hypoxia promotes expression of this differentiation marker even as it suppresses the overall EVT differentiation program.

Analysis of known Transcription Factor (TF) targets appropriately indicated that HIF1α was the most enriched TF associated with Cluster 1 (hypoxia) expression, while *GCM1* (Glial cells missing 1) was associated with high oxygen concentration (Figure S1I, S1J). *GCM1*, which is expressed in hTSCs, but upregulated upon differentiation (Figure S1K), was itself strongly downregulated in low oxygen conditions both at the RNA and protein level (Fig. 1J, 1K, S1L, Table S1). GATA3, reported to repress GCM1 by an indirect mechanism^26^, was upregulated in 2% O_2,_ but only slightly, suggesting another mechanism at work (Fig. S1H). To determine if GCM1 is regulated by the canonical hypoxia pathway, we used CRISPR interference (CRISPRi) to repress *VHL*, the ubiquitin ligase which targets HIF1α for destruction, in order to stabilize HIF1α at 20% O_2_. Repression of *VHL* in 20% O_2_ led to dramatic upregulation of *IGFBP3*, the most hypoxia responsive gene in hTSC and a known HIF1α target.^27^ We also observed downregulation of *GCM1* expression, confirming that *GCM1* is negatively regulated by canonical hypoxia response (Figure S1M).

### GCM1 is Essential for the Differentiation into Trophoblast Lineages

Since GCM1 is highly sensitive to oxygen concentration and is implicated in hTSC differentiation^26^, we generated GCM1-knockout (*GCM1^−/−^*Line 1) hTSC by deleting a small genomic region in exon 2 just after the ATG-start site to disrupt the translation of the DBD (DNA binding domain)^28^ and subsequent protein sequence (Fig. 2A). Unexpectedly, this deletion made an alternative splice site available, which spliced in before exon 3 (Fig. S2A). While this deletion still had the desired frameshift, we generated an additional line (*GCM1^−/−^* Line 2) deleting the entirety of exon 3 (Fig. 2A, S2A). hTSCs electroporated with a non-targeting (NT) sgRNA showed ubiquitous nuclear expression of GCM1, with TEAD4 loss in the highest GCM1 expressing cells (Fig. 2B), while both *GCM1^−/−^* lines showed loss of specific GCM1 signal (Fig. 2B). Consistent with recent reports^26,29,30^, *GCM1^−/−^*hTSC failed to differentiate to EVT or STBs, a result demonstrated by flow cytometry, immunofluorescent staining and RNA-seq of control and *GCM1*^−/−^ cells (Fig. 2C-2G, Figure S2B-F). *GCM1^−/−^* hTSCs showed lower expression of EVT and STB-specific genes and substantially failed to upregulate these genes upon directed differentiation (Figure S2G, Table S3).

**Figure 2.**
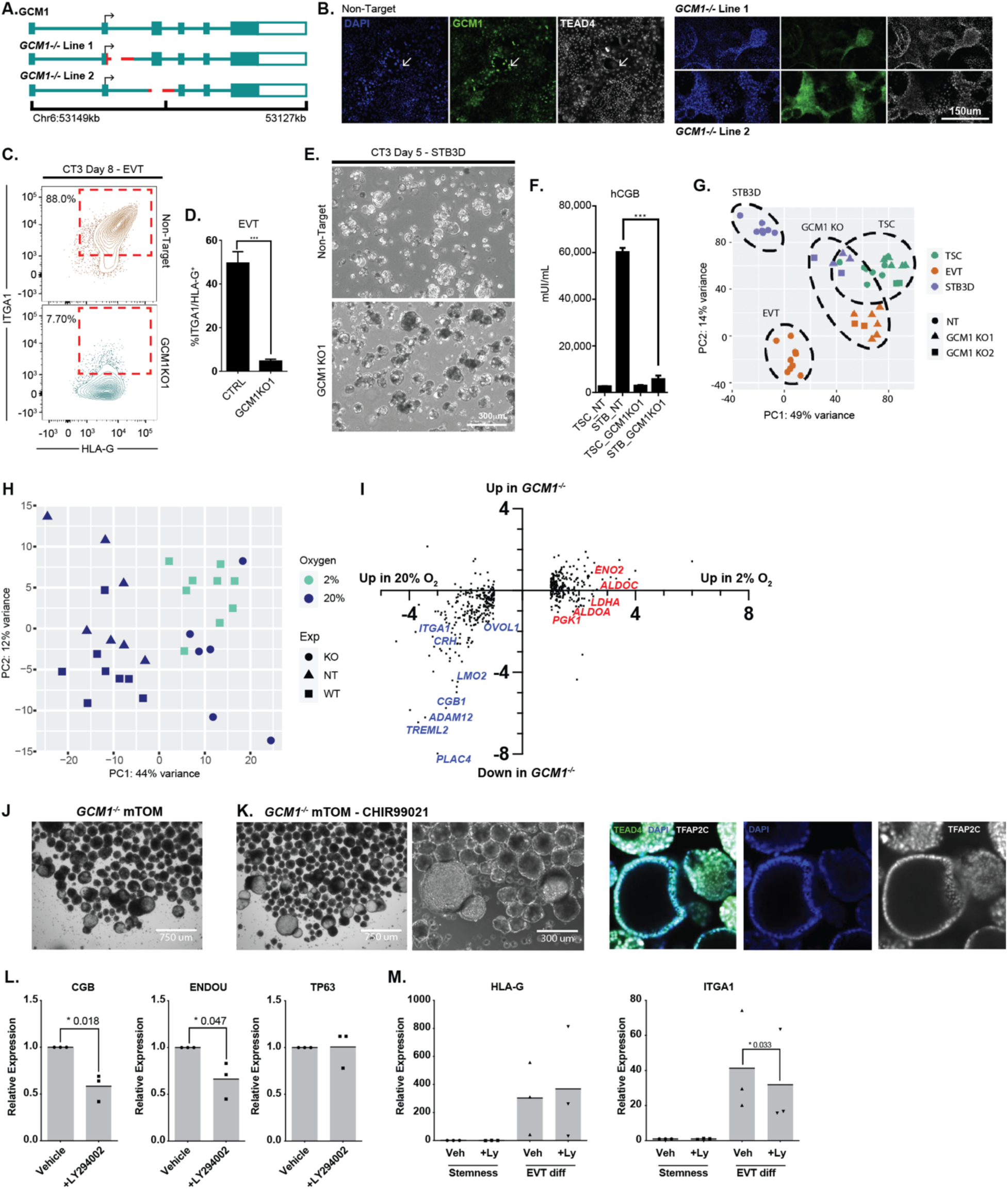
Impaired differentiation upon genetic or chemical reduction in GCM1 level. **A.** Strategies for mutation of GCM1 using a two sgRNA CRISPR approach. Lines were generated by deletion of the exon2/intron2 boundary, and by ablation of exon 3, either of which should disrupt the DBD of GCM1. **B.** Immunofluorescence staining of GCM1 and TEAD4 in control (Non-target, NT sgRNA) and *GCM1^−/−^* hTSC. Sporadic GCM1^+^ TEAD4^lo^ hTSCs are present only in NT control hTSCs. **C.** Flow cytometric analysis from EVT differentiation of *GCM1^−/−^* Line 1 and NT control TSC. NT hTSC differentiation produced ITGA1^hi^/HLA-G^hi^ cells whereas *GCM1^−/−^* TSC did not. **D.** Bar graphs showing formation of ITGA1^hi^/HLA-G^hi^ population from control and *GCM1^−/−^* TSC (n=5 replicates). **E.** 3D-STB formation of NT and *GCM1^−/−^* hTSC. Control hTSCs form a fluid-filled syncytium while *GCM1^−/−^* form a cluster of cells. **F.** hCGB ELISA was performed using supernatant from *GCM1^−/−^* and control hTSC (n=5 replicates). **G.** PCA comparing NT and *GCM1*^−/−^ hTSC, EVT, and STB3D. Note that *GCM1^−/−^* cells regardless of differentiation state cluster closer to the hTSC population, and similarity of *GCM1^−/−^* lines 1 and 2 (n=8 replicates for EVT, n= for STB3D). **H.** PCA of control (NT) and *GCM1^−/−^* hTSCs, compared with WT hTSCs grown at different O_2_ concentrations. Note that *GCM1^−/−^* hTSCs cluster on PC axis 1 with WT hTSCs grown at 2% O_2_. **I.** Scatterplot of genes differentially regulated in hypoxia (same set as Figure 1J) showing their relative expression in 2% and 20% O_2_ and their relative expression in *GCM1^−/−^*hTSC and control cells. Examples of placental differentiation genes are shown in blue, while genes involved in glycolysis are shown in red. **J.** Brightfield images of *GCM1^−/−^*TB-ORG cultured in mTOM media. **K.** (left) Brightfield images of *GCM1^−/−^*TB-ORG culture in mTOM media-CHIR99021 (right) Immunofluorescent staining for trophoblast markrers in *GCM1^−/−^* TB-ORG. **K.** Expression of genes associated with differnetiation (*CGB*, *ENDOU*) or stemness (*TP63*), normalized to the housekeeping gene *TBP*, in steady state conditions with 5μM LY294002 inhibitor or vehicle control (n=3 replicates). **L.** Expression of EVT genes upon differntiation to EVT with 5μM LY294002 inhibitor or vehicle control (n=3 replicates).

We then compared the effects of *GCM1* loss to the effects of hypoxia in hTSCs. While they did not cluster precisely together, *GCM1^−/−^* cells grown in 20% O_2_ showed similar positioning over principle component axis 1 with control hTSCs grown in 2% O_2_. This would indicate that a substantial portion of the differential gene expression associated with hypoxia is in fact a consequence of lower GCM1 level (Figure 2H). More specifically, we observe that a high proportion of genes downregulated in 2% O_2_, genes associated with trophoblast differentiation, are also downregulated in *GCM1^−/−^*. By contrast, genes upregulated in 2% O_2_ such as established HIF targets and factors that promote glycolysis^14^, are generally unaffected in *GCM1^−/−^* (Figure 2I).

Interesting morphological phenomena were observed for *GCM1^−/−^*hTSCs. When *GCM1^−/−^* lines were grown to over-confluency in hTSC media (TSCM), we observed three-dimensional dome-like projections. Much larger domes were observed in mTOM-C EVT-precursor media (see methods) (Figure S2H). We then cultured the *GCM1^−/−^* hTSCs using a modified form of a trophoblast organoid (TB-ORG) culture system. When cultured with standard mTOM media in in micro-V shaped wells with Matrigel omitted to allow free-floating organoids, the *GCM1^−/−^* TSCs formed hollow balls of cells (Figure 2J). Removal of CHIR99021 (mTOM-C) resulted in the further growth of these *GCM1^−/−^*shells (Figure 2K). At high cell number, the formation of branched, villous-tree like structures is observed (Figure S2I). Upon directed differentiation to EVT, these *GCM1^−/−^* TB-ORGs failed to enter an HLA-G^hi^ EpCAM^lo^ state and instead upregulated the surface marker ITGB6 (Figure S2J). ITGB6 is selectively present in column cytotrophoblasts^31,32^, the last GCM1^lo^ state before differentiation, suggesting failure to differentiate beyond this stage (Figure S2K-M).

TB-ORGs undergo spontaneous differentiation, forming a core of syncytiotrophoblast in the middle. We considered whether this could be prevented by reducing expression of GCM1. A published report in choriocarcinoma cells showed that hypoxia inhibits the PI3K/pAKT pathway, and that chemical inhibition of PI3K can lead to reduced expression of GCM1^33^. hTSCs treated with the PI3K inhibitor LY294002 showed some formation of dome-like structures akin to what is observed for *GCM1^−/−^* (Figure S2N, O). Likewise, LY294002-treated TB-ORG grown in mTOM-C conditions without Matrigel showed some propensity for formation of hollow cavities, though not to the same extent as *GCM1^−/−^* (Figure S2P). TB-ORG generated from primary CTBs and treated with LY294002 at standard steady state conditions with matrigel showed reduced expression of the STB markers *CGB* and *ENDOU* (Figure 2L) and a modest reduction in *ITGA1* expression upon EVT differentiation while HLA-G levels were not affected (Figure 2M), suggesting that chemical modulation of GCM1 level hampers certain steps of STB and EVT differentiation.

### GCM1 Positively regulates EVT and STB Specific Regulators

We performed Chromatin Immunoprecipitation Sequencing (ChIP-seq) for GCM1 from day 3 EVTs (CT3), a timepoint which we found was conducive to high quality ChIP data. We identified 2271 peaks with >4-fold enrichment over Input. Motif analysis of these sites showed extremely strong enrichment for the GCM-binding motif, with weaker enrichment for other TFs common in trophoblast (Fig. 3A). We confirmed enrichment at known GCM1 targets such as PGF and LMO2^29,34,35^ (Fig. 3B). We also conducted an assay for transposase accessible chromatin (ATAC-seq) to measure open chromatin regions in TSC, EVT and STB, which accorded well with published H3K27Ac ChIP-seq data from these cells (Fig. S3A). Comparison with the ATAC-seq data showed that GCM1 peaks corresponded to regions that show higher openness in EVT and STB than in TSC (Fig. 3C, Fig. S3B, Table S4). Furthermore, we used RAD (Region Associated DEG) analysis^36^ to correlate proximity of a GCM1 peak to the TSS of genes dysregulated upon GCM1 knockout. Genes downregulated in *GCM1^−/−^* cells were found in proximity to GCM1 ChIPseq peaks (Fig. 3D). Putative direct targets, genes proximal to GCM1 peaks and downregulated in *GCM1^−/−^* cells, include known placenta differentiation factors such as *PGF, LMO2, CGA, OVOL1, SYDE1*, and a range of *CGB* genes (Figure S3C).

**Figure 3.**
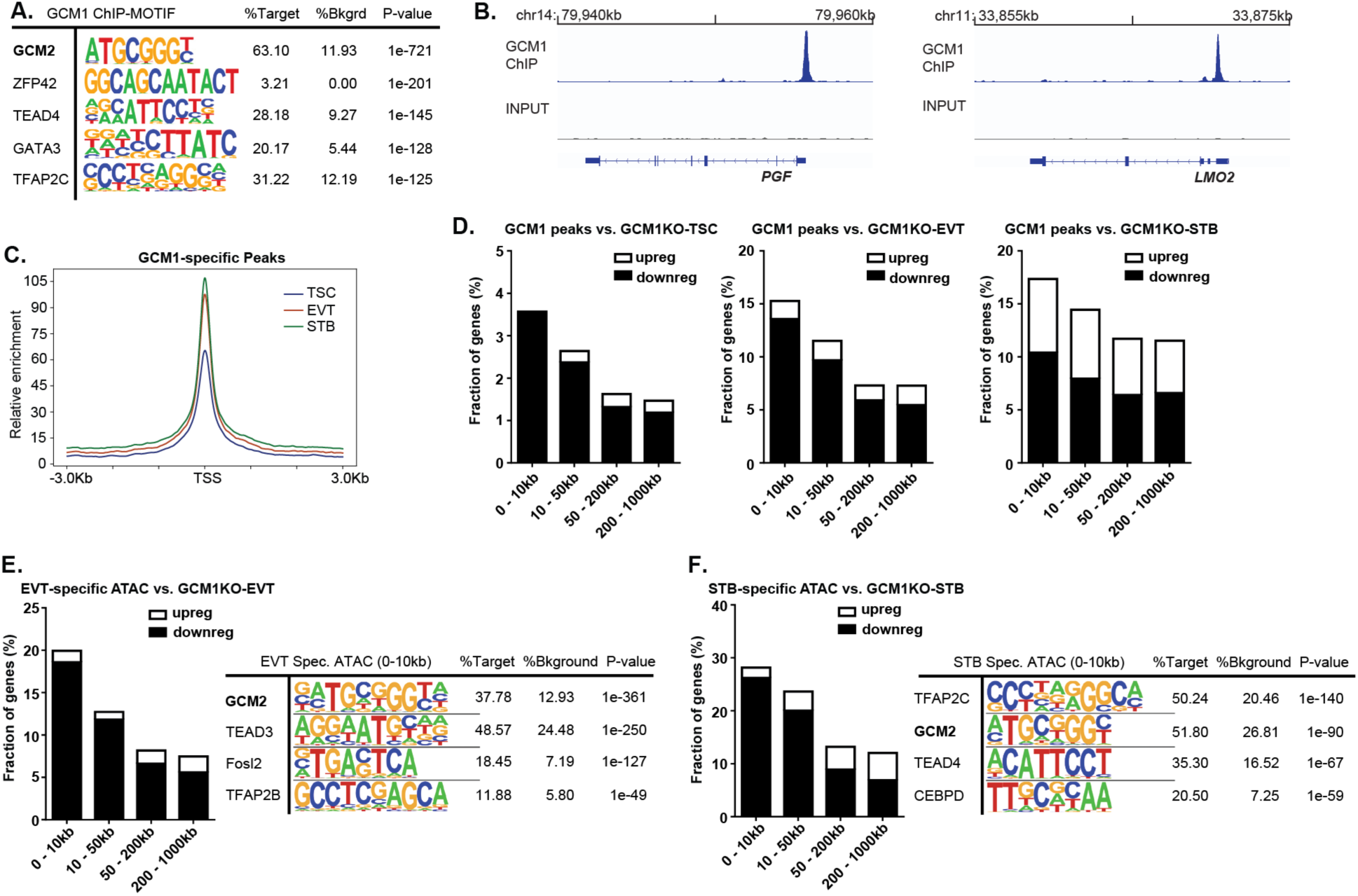
GCM1 positively regulates differentiation-associated genes. **A.** Motif analysis of GCM1 binding sites shows very strong enrichment for GCM motif, indicating successful and specific ChIP. **B.** GCM1 enrichment over *PGF* (left) and *LMO2* (right). **C.** ATAC-seq enrichment in TSC, EVT and STB over GCM1 binding sites. **D.** Plot showing the percentage of genes whose promoters are within a given distance of a GCM1 binding site that show upregulation or downregulation in *GCM1^−/−^*cells. **E.** Plot showing the percentage of genes whose promoters are within a given distance of an EVT-specific ATAC-seq site that show upregulation or downregulation in *GCM1^−/−^* cells (left), motif analysis for EVT-specific peaks (right) **F.** Plot showing the percentage of genes whose promoters are within a given distance of an STB-specific ATAC-seq site that show upregulation or downregulation in *GCM1^−/−^* cells (left), motif analysis for STB-specific peaks (right).

Further supporting the role of GCM1 in regulating differentiation, ATAC-seq analysis of TSC, EVT, and STB data showed enrichment of the GCM1 motif in EVT and STB-specific regions of open chromatin (Fig. 3E, 3F, 3SD). We also observed association of EVT and STB-specific ATAC-seq peaks with genes downregulated in *GCM1^−/−^* cells of the corresponding cell types (Fig. 3E, 3F).

### GCM1 Regulates CDKN1C and Trophoblast Overgrowth

One of the strongest GCM1 enrichment sites in the genome was found within the 11p15.5 imprinted locus^37^, as were several smaller peaks (Fig. 4A). This locus includes the transcripts *KCNQ1, KCNQ1OT1* and the protein CDKN1C (P57^kip2^), an inhibitor of cell division which binds to cyclin/CDK and blocks cell division^38^. Analysis of HiC data, which shows three-dimensional interaction of regions of chromatin, shows a high degree of interaction between the strong GCM1 binding site in *KCNQ1* and the promoter of *CDKN1C* (Fig. 4A, B)

**Figure 4.**
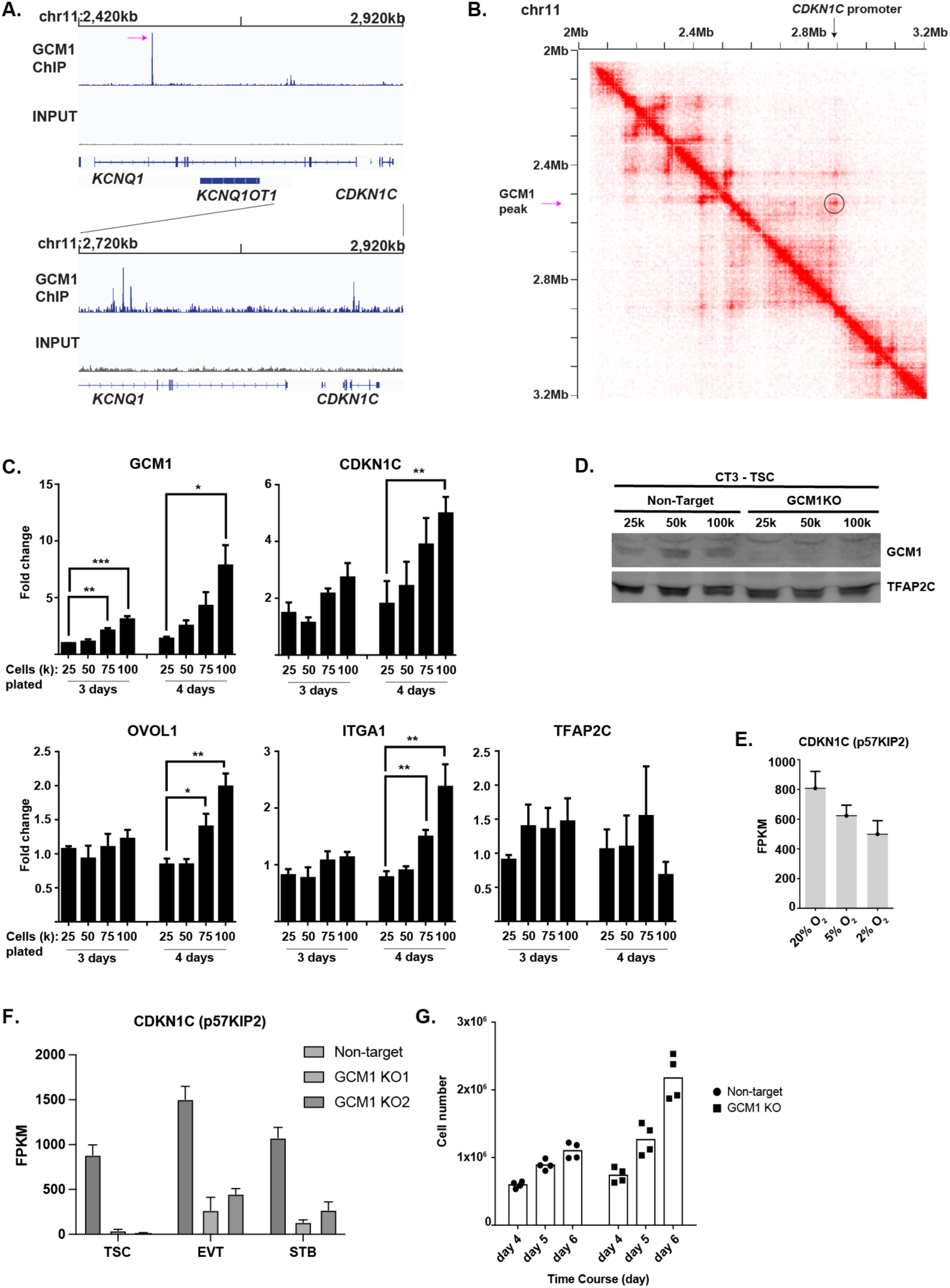
GCM1 positively regulates *CDKN1C* and contact inhibition. **A.** GCM1 enrichment over imprinted *KCNQ* locus. **B.** HiC interaction data over the *GCM1* locus. Note physical association between GCM1 binding site and *CDKN1C* promoter. In **B** and **C**, the highest GCM1 peaks is indicated with a magenta arrow. **C.** Expression of genes indicated in plating conditions (cell number and growth time) indicated. Note that plating at higher densities leads to higher expression of *GCM1* and *CDKN1C*. **D.** GCM1 protein levels increase with higher confluence. **E.** Expression of *CDKN1C* in oxygen concentration indicated. **F.** Expression of *CDKN1C* in control and *GCM1^−/−^* Line 1 and Line 2 hTSC and differentiated cells. **G.** Cell number after plating 50k cells and allowing cells to grow for indicated number of days. Note a leveling off in non-targeting cells as cell lines reach confluence, but continued growth in *GCM1^−/−^* hTSCs.

CDKN1C shows preferential expression from the maternal allele.^39^ A pregnancy abnormality called a full hydatidiform mole arises from an androgenetic pregnancy in which all genetic material is of paternal origin, and thus placental cells in hydatidiform moles feature loss of *CDKN1C* as well as persistent trophoblastic outgrowth.^40^ hTSC upregulate *CDKN1C* at high confluence, and *CDKN1C^−/−^* hTSC lose contact inhibition and continue growing after reaching confluence.^38^

When we grew hTSCs to high confluence, we observed upregulation of *GCM1* and *CDKN1C* in tandem (Fig. 4C,D), with more modest increases in other differentiation markers (Fig. 4C). We also observed reduced *CDKN1C* expression in hypoxia (Fig 4E), where levels of GCM1 are lower (Fig. 1I, S1L). Further consistent with direct regulation by GCM1, *CDKN1C* was dramatically downregulated in *GCM1^−/−^* hTSCs and differentiated cells (Fig. 4F). While control cells stopped dividing as confluence occurs, GCM1KO cells continued to expand, similar to the reported *CDKN1C^−/−^*phenotype (Fig, 4G).^38^ These results collectively indicate that GCM1 acts upstream of *CDKN1C* in response to confluence and controls its expression.

## Discussion

Considering the positive effect of hypoxia on spongiotrophoblast differentiation in mouse, how can we explain the generally inhibitory effect of low oxygen on EVT differentiation in humans? A critical difference between mice and humans, as illustrated in our work and others^26,29,30^, is that GCM1 is essential for both EVT and STB formation in human but only STB formation in mouse. Hence, if hypoxia negatively regulates GCM1 in both mice and humans, this would be predicted to have an inhibitory effect on EVT differentiation in humans but not on spongiotrophoblast or trophoblast giant cell (TGC) formation in mice. We also find, somewhat counterintuitively, that hypoxia has a positive effect on HLA-G expression even though it has an overall negative effect on EVT differentiation. It is worth noting here that hTSCs, while clearly bipotent, express some markers consistent with cell column cytotrophoblast, indicating that they may be more EVT-like than typical villous cytotrophoblasts^41,42^. Hence, the observation that hypoxia promotes proximal-column EVT transcriptional program^22^ is not necessarily incompatible with a role in hTSC self-renewal.

With regard to pathology, preeclampsia is widely understood to feature inadequate remodeling of maternal arterioles and concomitant reduced blood and oxygen availability for the placenta^43^, and higher levels of HIF-1α and/or HIF-2α protein have been observed in preeclamptic placenta^44^. Excess of undifferentiated cytotrophoblasts, which our study predicts would result from hypoxia, has also been reported^45^.

TSC-organoids derived from human placenta collected over several gestational stages have the consistent feature that STB differentiation occurs inside the organoid.^46–49^ This may reflect higher pressure inside the organoid, or lack of access to EGF, but this organization is inverted with respect to the bilayer formation of chorionic villus.^50^ Two groups have succeeded in finding conditions in which STBs form on the outside of the organoid, though the resulting organoids cannot be propagated^51,52^. We reasoned that reduction of *GCM1* level, as reported in literature from the PI3K inhibitor treatment, could allow sustained culture of undifferentiated organoids. We did observe reduced, though not eliminated, spontaneous differentiation. Interestingly, the published^33^ mechanism by which LY294002 reduces GCM1 expression, via activation of GSK3β which then phosphorylates and degrades GCM1, would not be expected to work in TSCM media conditions in which GSK3β is perpetually inhibited via treatment with CHIR99021. Hence, it is something of a mystery how LY294002 prevents spontaneous differentiation in the organoid model, and future research in this area could yield improved culture conditions.

In a CRISPR dropout screen of hTSCs, *GCM1* is a growth-restricting gene, whose deletion promotes cell growth^53^. This is likely due to two mechanisms. As shown above, hTSCs, especially at high confluence, undergo some spontaneous differentiation which is suppressed by loss of GCM1. Furthermore, *GCM1^−/−^* hTSCs show greatly reduced *CDKN1C* expression, limiting contact inhibition. Indeed, *CDKN1C* is also a growth-restricting gene in CRISPR screens^30,53^. Interestingly, despite its growth-restricting properties in culture, *GCM1* is not a known tumour suppressor in gestational choriocarcinoma (GC)^54–56^. In the case of hydatidiform-mole derived GC, *CDKN1C* expression is already lost, but GC can also arise from non-molar pregnancy and there is no evidence of GCM1 mutation in these diseases either. This may be because while loss of *GCM1* causes loss of contact inhibition and uncontrolled growth, it also precludes EMT and invasiveness. At the same time, analysis of mutations and karyotypic abnormalities in GC remains limited and there is almost nothing known about the mutational profile of the related placental cancers, placental site trophoblastic tumor (PSTT) and epithelioid trophoblastic tumor (ETT)^57–59^. New roles for GCM1 in placental development and placental cancer may as yet be discovered.

### Experimental Procedures

#### Cell Culture-Maintenance of hTSC

CT1 and CT3 lines were generously provided from Dr. Arima’s lab in Japan. TSC were cultured in TSC basal media containing DMEM F-12 (GIBCO), 1x ITS-X (GIBCO), 0.3% BSA (WISENT), 1% Penicillin/Streptomycin (GIBCO), 1% ESC qualified Fetal Bovine Serum (GIBCO), 0.1mM β-mercaptoethanol (GIBCO), 15μg/ml L-ascorbic acid (SIGMA), and 50ng/ml recombinant hEGF (GIBCO). To this, made fresh, 0.75mM valporic acid, 2μM CHIR99021 (Cayman Chem), 0.5μM A8301 (Cayman Chem), 1μM SB431542 (Cayman Chem), and 5μM Y27632 (Cayman Chem) was added to make TSC Media (TSCM). TSCs were dissociated using TRYPLE (GIBCO) diluted with PBS to 30% and incubated at 37°C for 10 min. Deactivation of TRYPLE was done by using 1:1 vol of 0.5mg/ml Soybean trypsin inhibitor (GIBCO) diluted in PBS. For experiments performed in Figure 1, hTSCs were passaged on to 5μg/ml of Collagen IV (Corning) coated plates, but due to product availability, TSC from experiments in Figures 2-4 were passaged onto Laminin 511(SIGMA) coated plates.

#### Cell Culture-Differentiation of hTSC to EVT

TSCs were converted to EVTs using a modified TSC basal media containing DMEM F-12 (GIBCO), 1x ITS-X (GIBCO), 0.3% BSA (WISENT), 1% Penicillin/Streptomycin, and 0.1mM β-mercaptoethanol (GIBCO). For days 1-2, TSCs were passage onto 1μg/ml Collagen IV coated plates in mod. TSC basal with added 5% Knockout-serum replacement (KSR) (GIBCO), 5μM Y27632 (Cayman Chem), 3uM A8301 (Cayman Chem), 100ng/ml NRG1 (NEB) and 2% GFR-Matrigel (Corning) while the media is still cold (EVTM). Days 3-5, the media is changed to ETVM without NRG1 and reduction to 0.5% GFR-Matrigel. Days 6-8, the media is changed to the previously mentioned ETVM but this time without NRG1, KSR, and reduction to 0.5% GFR-Matrigel. At the end of day 8, differentiated EVTs can be assessed by Flow cytometry.

#### Cell Culture-Differentiation of hTSC to STB3D

TSCs were converted to STBs using a modified TSC basal media containing DMEM F-12 (GIBCO), 1x ITS-X (GIBCO), 0.3% BSA (WISENT), 1% Penicillin/Streptomycin, and 0.1mM β-mercaptoethanol (GIBCO). For days 1-2, TSCs were passage on to suspension plate (Sarsted) in TSC basal media containing 5μM Y27632, 2μM Forskolin (Cayman Chem), 5% KSR (GIBCO), and 50ng/ml recombinant hEGF (STBM). On day 3, cell clusters were collected and pulsed-spin for 30sec to separate single cells from STB fusion-clusters and replated in STBM. At day 5, STB-3D are ready to be assessed by immunofluorescence and/or hCG-ELISA

#### Trophoblast organoid and EVT differentiation (modified protocol without Matrigel)

3D- TSCs (TB-ORG) of control and *GCM1^−/−^* cells were cultured based on a modification from the Okae *et al*. 2018 and Haider *et al*. 2022 publications. Our modified-TOM (mTOM) media contains DMEM F-12 (GIBCO), 10mM Hepes (GIBCO), 1x ITS-X (GIBCO), 2mM GlutaMax (GIBCO), 1x Penicillin/Streptomycin (GIBCO), 0.2% ESC-FBS (GIBCO), 50ng/ml rhEGF (Invitrogen), 3μM CHIR99021 (Caymen Chem.), 2μM A8301 (Caymen Chem.), and 5μM Y27632 (Cayman Chem.) Instead of Matrigel embedding, micro-V-shape wells (AggreWell 400, Stemcell Tech.) were used to generate organoids. ∼50,000 cells were resuspended in 1ml mTOM per 24 well and centrifuged in a plate spinner for 5 min at 1000 rpm to collect cells to the bottom at ∼100 cells per V-shape well. After 24 hours, 500μl of media are gently removed and either mTOM is replaced or mTOM without CHIR99021 (mTOM-C) for CTB-CCC/precursor to EVT formation was replaced every other day over a ∼10-day period. For LY294002 treat, treatment up to 14 days may be required to physically see by microscopy cavity formation.

#### Cell culture – standard maintenance and EVT differentiation of trophoblast organoids

TB-ORG were generated and cultured according to a recent publication^60^. Briefly, villous cytotrophoblasts (vCTBs) were isolated (6 – 7^th^ week of gestation, n=3), and 1 × 10^5^ cells were embedded in Matrigel and cultivated in advanced DMEM/F12 (Invitrogen) supplemented with culture medium containing 10 M HEPES, 1 x B27 (Gibco), 1 x ITS-X (Gibco), 2mM glutamine (Gibco), 0.05 mg/ml gentamicin (Gibco), 1 µM A8301 (R&D Systems), 50 ng/ml recombinant human epidermal growth factor (rhEGF, R&D Systems), 3 µM CHIR99021, and 5 µM ROCKi (Y27632, Santa Cruz). After the first passaging, ROCKi omitted. The medium was changed every 2 – 4 days, and TB-ORG were split after 5 – 7 days. To test effects of LY294002 under stemness conditions, TB-ORG of passage 2 were treated with DMSO (vehicle) or 5 µM LY294002 for 10 days. Culture media were changed every 2 – 3 days.

For EVT differentiation, passage 2 TB-ORG were incubated with TB-ORG medium lacking CHIR99021 (TB-ORG-DIFF). To test effects of LY294002 on EVT lineage formation and differentiation, TB-ORG were incubated in TB-ORG-DIFF medium supplemented with DMSO (vehicle) or 5 µM LY294002 for 10 days. EVT formation was monitored and bright field images were taken every 2 – 3 days.

### Confluence experiment

CT1 hTSCs were plated in 24 well plates at 4 different densities (25, 50, 75 and 100K cells), in duplicates, and incubate for 48 or 72 hours in order to reach 100% confluence at the highest density. Cells were collected using TRYPLE 30% at 37°C for 10 min followed by addition of trypsin inhibitor. Single cells obtained were centrifuged to obtain a pellet, washed with PBS and flash frozen and stored at −80°C.

#### Generation of CRISPR Knockout lines by Nucleofection

CRISPR guides were designed using IDT-Custom Alt-R^™^ CRISPR-Cas9 guide RNA program. sgRNA guides targeting a small region of nucleotide just after the ATG-start site in exon 2 and introns 2-3 (sgRNA1-UCU UCA GAA UCA AAG UCG UC and sgRNA2-ACU AUU AAC AUG CGG AGA CC). Additional deletion of exon 3 were designed (sgRNA3-GAG CGC UGC UCA GAU AGC GA and sgRNA4-AGA CCU AAG AGC AAU CAG UG). Cas9-sgRNA ribonucleoprotein complexes were generated from sgRNA and Cas9 protein (Synthego). In brief, 75pmol of each individual sgRNA is complexed with 10pmol of Cas9 protein in Cas9 annealing buffer (NEB) for 10min. In the meantime, TSCs are dissociated with 30% TRYPLE and reconstituted in PBS to a concentration of 1 × 10^5^ cell/μl. Pre-complexed RNPs sgRNA1 and sgRNA2 are combined together with 10μl of cell solution and 20μl of P3 solution (LONZA), and transferred to a cuvette to be nucleofected using an Amaxa 4D nucleofector (Lonza) with pulse code CA137. Immediately after, 150μl of TSC media is added to the cuvette to neutralize the reaction and transfer cells to a freshly prepared 10cm plate coated with Lam-511 and TSCM for generating single clones. 2mg/ml Collagenase V solution was used dissociate colonies in order to pick single clones. Deletions were confirmed by genotyping (GCM1KO1: Forward-TTGTATGAGGACTTGTGCATAACAA, Reverse-GCCATTGGTTACAGATGACAAC, GCM1KO2: Forward-ATGGAACTCACAGGGGCTAT and Reverse-TAACAGGAGCCTTCAGTCCA).

#### Chromatin immunoprecipitation

Cell samples are fixed with 0.66% paraformaldehyde (FisherSci) diluted with PBS and is quenched by adding glycine to a final concentration of 0.125M. Nuclear lysis extraction begins with nuclear lysis buffer (50mM HEPES pH 7.8, 0.5% Triton X-100, 1mM EDTA, 0.5mM EGTA, 140 mM NaCl, 10% glycerol and 1% NP-40). Nuclei are resuspended in nuclear wash buffer (10mM Tris-HCl pH 8.0, 200mM NaCl, 1mM EDTA, and 0.5mM EGTA). Nuclear pellets are then resuspended in SDS lysis buffer (50mM Tris-HCl pH 8, 10mM EDTA, 1% SDS, and 1% Triton X-100). Nuclei samples are transferred to a 1ml tube (Bioruptor) and keep cold before sonication. Sonication was performed using the Diagenode Bioruptor sonicator (settings: 30sec on, 30 sec off, 30-35 cycles). Then diluted with Dilution buffer (25mM Tris-HCl pH 8, 150 mM NaCl, 3mM EDTA, and 1% Triton X-100). A pre-clearing step is performed by using 40μl of pre-washed Protein G Sepharose beads (SIGMA, P3296-5ml) combined sonicated sample. 2µg of GCM1(HPA001343) antibody is added to each pre-cleared sample and then rotated overnight at 4°C. Pre-washed Protein G Sepharose beads are added to the ChIP samples and washed with three buffers: DB150 (25mM Tris-HCl pH 8.0, 150mM NaCl, 3mM EDTA, 1% Triton X-100, and 0.05% SDS), DB500 (25mM Tris-HCl pH 8.0, 500mM NaCl, 3mM EDTA, 1% Triton X-100, and 0.05% SDS), Buffer III (10mM Tris-HCl pH 8.0, 250mM LiCl, 1% Sodium Deoxycholate, 1% NP-40, and 1mM EDTA, and lastly TE buffer (10mM Tris-HCl pH 8.0, 1mM EDTA). DNA is eluted with 200µl elution buffer (100 mM NaHCO3, and 1% SDS) and incubated at 65°C overnight for decrosslinking. Ethanol precipitation is performed to collect ChIP material, and further purification was performed with Geneaid Gel/PCR cleanup protocol. Purified DNA fragments passed to sequencing library preparation.

#### ATAC-Seq

ATAC-seq library preparation was performed on 1 × 10^6^ freshly cultured cells using a commercially available ATAC-Seq kit from Active Motif (#53150, Carlsbad, CA). We followed manufacturers protocol with some minor modification. Each sample was lysed in the ATAC-seq lysis buffer. Next, the samples were processed for the transposase reaction. After cleanup of the transposed DNA, samples were stored at −20 °C until library amplification. Samples were subsequently thawed at room temperature and library construction completed according to manufacturer protocol. Libraries were quantified using a Qubit dsDNA BR Assay Kit (Q32853, Thermo-Fisher) and the size was determined with a High Sensitivity DNA Bioanalyzer Kit (5067-4626, Agilent, Santa Clara, CA) and sequenced on a NovaSeq 6000 (Illumina, San Diego, CA) using Nextera Sequencing primers.

#### RNA isolation and qPCR

Total RNA isolation used manufacturing protocol indicated by Sigma-Aldrich RNAzol^RT^ R4533. Qubit™ RNA BR Assay kit (Q10211) was used to measure RNA concentration.

First-strand cDNA was generated using the SensiFast cDNA Synthesis kit (Froggabio). Quantitative PCR was done using PowerUP SYBR^TM^ Green PCR (Invitrogen, A25742) on Quantstudio 5 (Applied Biosystems) with the following cycling conditions: 50 °C 2 minutes, 95 °C 20 seconds, 45x (95 °C 3 seconds, 60 °C 30 seconds), 95 °C 1 second). The qPCR reaction was performed using 1x concentration of PowerUP SYBR Green Master Mix, 5ng of template mRNA and 0.5 µM of primer mix in a total of 7 µL reaction. The expression of target genes was normalized to the housekeeping gene Rab7a and the sequence of primer used for qPCR are provided in the table below.

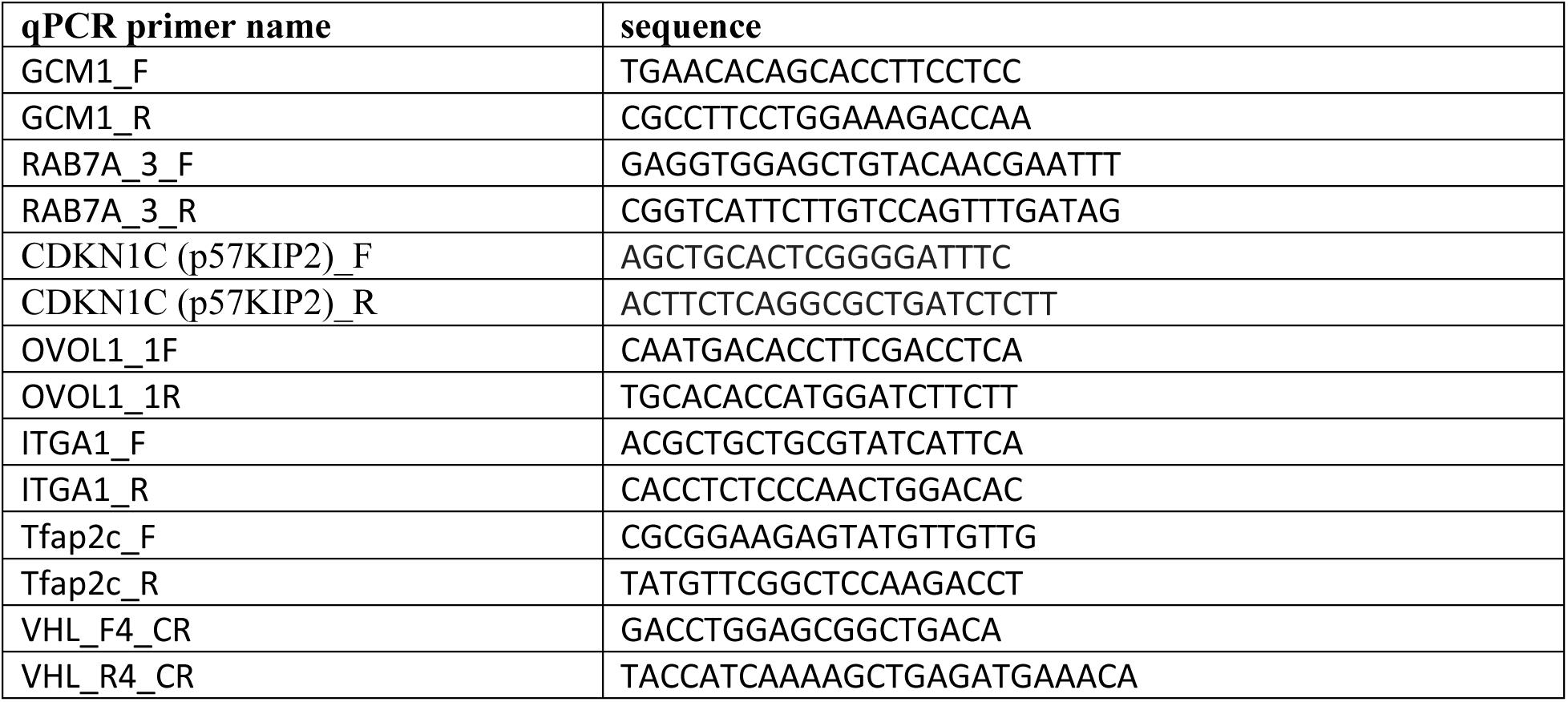

#### RNA isolation and qPCR from TB-ORG (Figures 2L,M)

TB-ORG were washed with ice-cold PBS and re-suspended with PeqGold Trifast (PeqLab). Homogenization of TB-ORG was supported using the Precellys 24 (CK-Mix tubes, 5000 rpm, 1 x 20 sec, PeqLab) and RNA isolation was performed as indicated by the manufacturer. RNA (1 µg per sample) was reverse transcribed (RevertAid H Minus Reverse Transcriptase, Thermo Scientific) and qPCR was performed (7500 Fast Real-time PCR system, Applied Biosystems). The following TaqMan Gene Expression Assays (ABI) were used: *CGB* (Hs00361224_g), *ENDOU* (Hs00195731_m1), *TP63* (Hs00978340), *HLA-G* (Hs00365950_g1), and *ITGA1* (Hs00235006_m1). Signals (ΔCt) were normalized to TATA-box binding protein (*TBP*, 4333769F).

#### Library preparation

For RNA library synthesis, in brief, purified mRNA was cleaned using the NEBNext Poly(A) mRNA Magnetic Isolation Module kit (NEB E7490) and final library generation was created with the NEBNext® Ultra RNA Library Prep Kit for Illumina® (NEB E7530) following manufacturing instructions. Barcoding came from TruSeq Unique Dual Indexes (Illumina, San Diego, CA). Qubit™ 1x dsDNA HS Assay kit (Q33231) was used to measure synthesis of the library.

#### Sequencing Analysis

For RNA and ChIP/ATAC sequencing analysis, Genpipes 4.3.2 (https://bitbucket.org/mugqic/genpipes) provided the pipeline for basic sequencing processing. For RNA-seq, sequencing quality and adaptor removal was trimmed with trimmomatic (v0.36). Trimmed fastq files alignment was performed with STAR aligner (v2.7.8a) using hg38/GRCh38 (ensembl v104). Picard (v2.9.0) then used merged, marked duplicates, identifed unique read, and sorted .bam files proceeding alignment. Read counts were collected using HTseq-count and StringTie (v1.3.5). Differentially expressed gene (DEG) comparison was performed with DESeq2 package on RStudio (R v4.3.1). Correlation matrix, hierarchal gene cluster analysis, and PCA were generated in R. Gene pathway analysis were performed on ConsensusPathwayDB and EnrichR. For ChIPSeq and ATACseq, in brief, raw fastq files trimmed using trimmomatic (v.0.36). Then qualified fastq reads were mapped with BWA (v.0.7.17) with post processing with sambamda (v0.8.1) to merge replicates, mark and filter duplicates, and remove blacklist regions. Peak calling and differential-bind was performed with MACS2 (v2.2.7.1). Gene annotations and motif analysis was performed with Homer (v4.11). Bam to bigwig bamCoverage from deepTools (v3.5.1) was used to generate the tracks for viewing on IGV (v2.9.4).

#### hCG ELISA

hCG secretion was measured using an hCG AccuBind ELISA (Monobind) according to manufacturer instructions.

#### Flow cytometry

Dissociated single cells were first washed with 1% KSR-PBS solution. Cell samples were incubated with fluorescently conjugated antibodies for 15min at room temperature. Post-incubation, the samples were washed once with 1% KSR-PBS, then resuspended in 1% KSR-PBS containing DAPI nuclear counterstain to identify live or dead cells. Data acquisition was preformed using the LSR Fortessa and data analysis was performed on FlowJo v10.

#### Western blot

Dissociated single cell samples were washed with cold PBS and lysed with Laemmli buffer without blue dye and boiled at 95°C for 5 minutes. Protein concentration was determined using a standard Bradford assay. Standard Bio-RAD SDS-PAGE system was used to separate proteins and transfer it to PVDF membrane (Millipore). Membrane blocking, primary and secondary antibody incubations are diluted in Odyssey Blocking buffer (LICOR). Infrared conjugated secondary antibodies were used for detection and visualization of the protein of interest on the membrane with the LICOR-Odyssey Imager.

#### Immunofluorescence

Glass coverslips were precoated with 5µg/ml Collagen IV (CORNING) overnight before cell attachment. Cells were grown for a determined amount of time for its corresponding experiment. Coverslips were fixed with 4% PFA for 20 min. at room temperature. PFA solution was washed 3 times with PBS before permeabilization with permeabilization buffer (PBS + 5% donkey serum + 0.1% Trition-X100) for 30 min. Primary and secondary antibodies were diluted in permeabilization buffer and incubated for 1-2 hours for each process. Coverslips were washed with PBS containing DAPI nuclear counterstain and mounted on glass slides using Pro Long Gold (Invitrogen). Imaging analysis was performed using the Axiovert (Zeiss).

## Supporting information

Supplementary Table Guide

## Data availability

Sequencing data was deposited to the GEO repository with the following Accession numbers. RNA-seq (GSE276594 and GSE276595), ATAC-seq (GSE276588), ChIP-seq(GSE276590).

## Author Contributions

J.K.C., S.Y.K., A.P., T.M., J.S., J.G., J.Z., P.D., M.J.J., conducted the experiments. J.K.C. and D.S. conducted the bioinformatic analysis. S.J.R., S.P., S.H., and W.A.P. supervised the experiments and analysis.

## Acknowledgments

We thank the Rosalind &Morris Goodman Cancer Research Center Flow Cytometry core, the SickKids Centre for Applied Genomics facility, the La Jolla Institute for Allergy and Immunology Sequencing Core, and the Canada Michael Smith Genome Sciences Center at BC Cancer for their dedicated service. This work was funded by the New Frontiers in Research Fund (NFRF) grant NFRFE-2018-00883 and the Canadian Institutes of Health Research (CIHR) project grant PJT-166169 to W.A.P., the NIH grants HD101319, HD062546 and HD103161 to S.P. and the Austrian Science Fund P34588-B, P36159-B to S.H. W.A.P. was supported by an FRQS Chercheurs-boursier. J.K.C. was supported by a Fonds de recherche Santé Québec graduate fellowship and studentships from the McGill University Faculty of Medicine.

---

**Figure S1.**
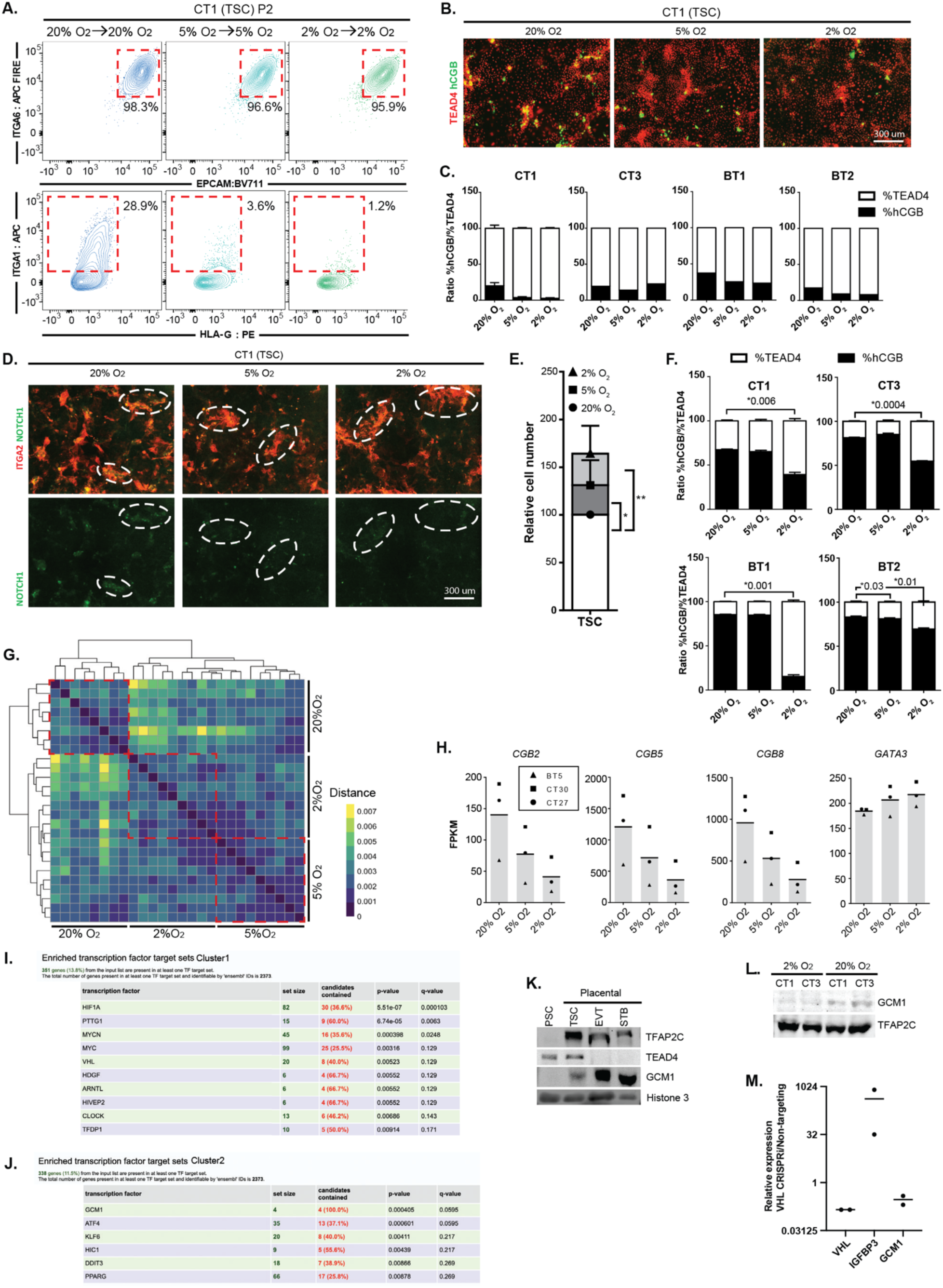
Reduced and impaired hTSC differentiation in hypoxic conditions. **A.** Trophoblast stem cells shown in Figure. 1A were passaged and cultured for additional 72hrs in varying levels of oxygen (20%, 5%, 2% O_2_). Continued low oxygen tension causes further reduction of ITGA1^+^ cell population. **B.** Spontaneous differentiation of hTSC to STB in as indicated by loss of TEAD4 and gain of hCGB. Note trend toward higher hCGB staining at 20% O_2_. **C.** Quantification of spontaneous hCGB expression across multiple cell lines at oxygen concentration indicated (4 cell lines, n=4 for each cell line). **D.** hTSC in the varying oxygen conditions were grown to over maximum confluency. At regions where overgrowth causes increase cell-to-cell contact and cell pile-up, spontaneous nuclear NOTCH1 signal is observed. In the low oxygen cultures, NOTCH1 expression is not detected. **E.** Relative numbers of TEAD4+ cells per unit area (4 cell lines, n=4 for each cell line). **F.** Quantification of spontaneous hCGB expression across multiple cell lines at oxygen concentration indicated (4 cell lines, n=4 for each cell line). **G.** Correlation matrix showing sample clustering of RNA-seq data from hTSCs in culture conditions indicated (3 cell lines, n=3 for each line in each condition). **H.** Bar graphs showing FPKM of specific genes of interest (n=3 replicates, except BT2 at 20% O_2_ n=2). **I., J.** ConsensusPathwayDB analysis of Cluster 1 and Cluster 2 from Figure 1H, identifying TF targets with significance from each cluster. **K.** Western blot comparing pluripotent stem cell (PSC) that don’t express placental markers, with hTSC and differentiated EVT and STB to highlight specific expression patterns. **L.** Western blot for GCM1 in 2 cell lines grown in 2% and 20% O_2_. TFAP2C is the loading control. **M.** Ratio of expression for genes indicated in hTSCs subjected to CRISPRi-targeted degradation of VHL relative to control hTSC. Note reduced expression of GCM1, while known hypoxia target IGFBP3 is dramatically upregulated.

**Figure S2.**
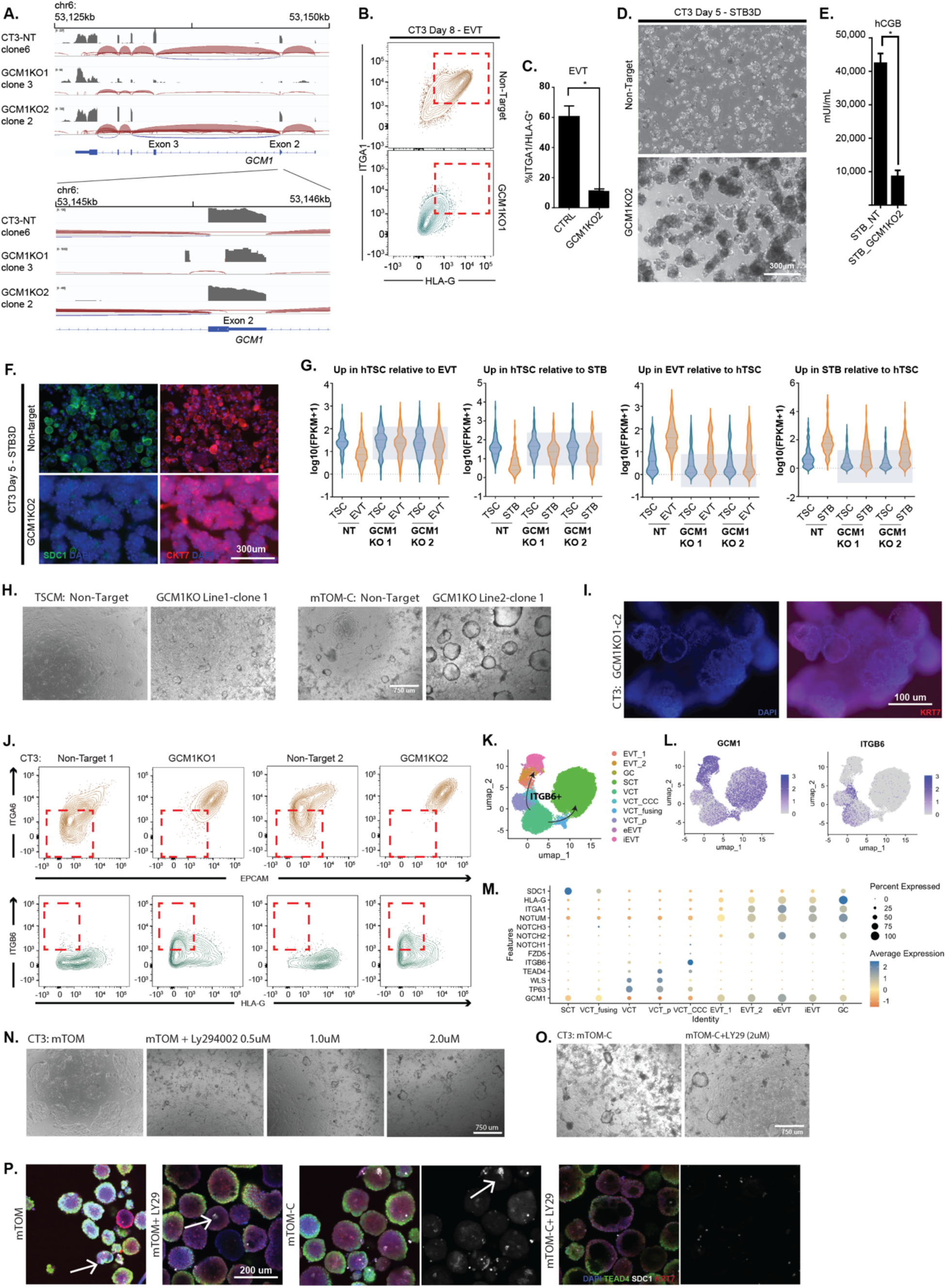
Impaired differentiation upon genetic or chemical reduction in GCM1 level. **A.** Sashimi plot across the genomic region of *GCM1*. Representative clones were chosen. Normal splicing is observed from non-target line. *GCM1^−/−^*Line1 (from right to left) show a deletion at the distal tip of exon 2 but an alternative slice site forming just after. *GCM1^−/−^* Line 2 shows the complete skipping of exon 3. **B.** Flow cytometric analysis from EVT differentiation of *GCM1^−/−^* Line 2 and NT hTSC. NT cells differentiation produce ITGA1^hi^/HLA-G^hi^ cells whereas *GCM1^−/−^*TSC do not. **C.** Bar graph showing formation of ITGA1^hi^/HLA-G^hi^ population from control and *GCM1^−/−^* Line 2 hTSC (n=3 replicates). **D.** STB3D formation of NT and *GCM1^−/−^* Line 2 hTSC. Control hTSCs form a fluid-filled syncytium while *GCM1^−/−^*form a cluster of cells. **E.** hCGB ELISA was perform using supernatant from *GCM1^−/−^* and control hTSC (n=3 replicates). **F.** Control and *GCM1^−/−^* Line 2 STB3D stained for the STB-marker SDC1 and the pan-placental marker CKT7. Note absence of SDC1 in *GCM1^−/−^*. **G.** Violin plot showing expression of genes specific to hTSC, EVT or STB in cell types indicated. Differentiated *GCM1^−/−^* cells fail to express differentiation markers and retain expression of TSC markers instead. **H.** *GCM1^−/−^*-hTSC were grown in mTOM with and without CHIR99021. Dome-like projections appeared in regions of high cell density. **I.** Immunofluorescent staining of *GCM1-/-* trophoblast organoids. **J.** Flow cytometry of NT and *GCM1^−/−^*-3D-TSC differentiated to EVT. *GCM1^−/−^* hTSCs fail to upregulate the EVT marker HLA-G but do upregulate the cell-column marker, ITGB6. **K.-M** Reanalysis of Arut. *et al.* 2022 single RNA-seq profiling the several subtypes found in early villus of the placenta. **K.** Cell types in placenta, with path of differentiation indicated. (GC=Giant Cell, VCT = villous CTB, VCT_p=proliferating CTB, VCT_CCC = cell column cytotrophoblast eEVT=endovascular EVT, iEVT= interstitial EVT). **L.** Expression of GCM1 and ITGB6 in cells shown in (**K**). **M.** Expression of genes indicated in cell types indicated. Note that ITGB6 is associated with cell column cytotrophoblast. **N.** hTSC cultured in mTOM media with varying quantities of LY294002. **O.** hTSC cultured in mTOM-C with or without LY294002 2µM. **P.** day14 TB-ORG grown in mTOM or mTOM-C with or without LY294002 2µM treatment and IF stained for DAPI, TEAD4, SDC1, and KRT7. Arrows mark areas of SDC1expression.

**Figure S3.**
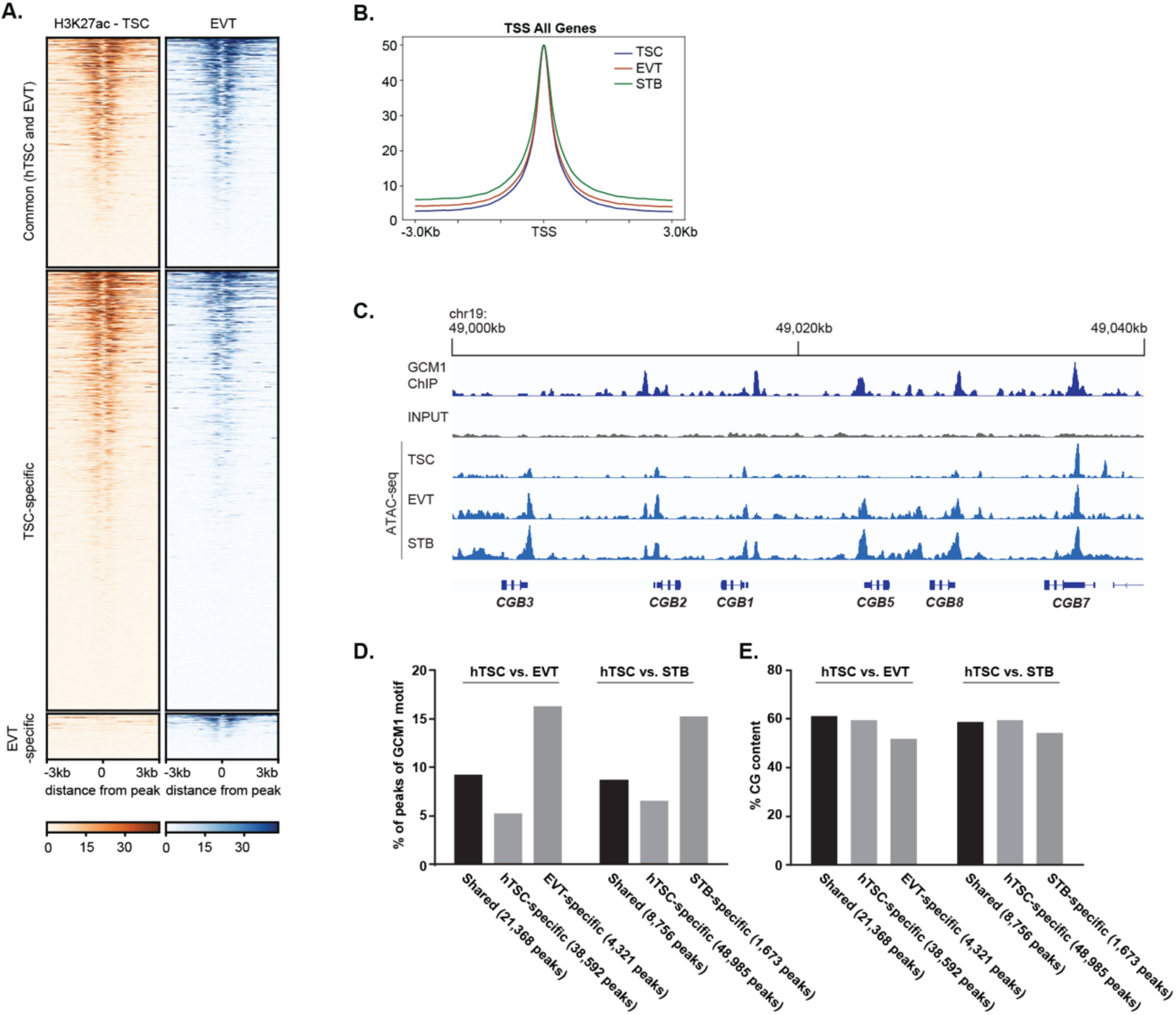
GCM1 positively regulates differentiation-associated genes. **A.** Heatmap of H3K27Ac enrichment over common, hTSC-specific, and EVT-specific ATAC-seq peaks in hTSC and EVT. Note correspondence of H3K27Ac enrichment with ATAC enrichment in each set. **B.** Metaplot of ATAC-seq data from TSC, EVT and STB over all gene TSS after normalization. **C.** GCM1 ChIP-seq and ATAC-seq data plotted over the *CGB* locus. **D.** Percentage of peaks in each category containing GCM motifs. **E.** GC content of ATAC-seq peaks in each category. GCM1 has a GC-rich motif, but the higher frequency of GCM1 sites observed in **(D**) cannot be explained by difference in underlying GC richness.

