## Supplementary Table Guide for "Hypoxia and loss of *GCM1* expression prevents differentiation and contact inhibition in human trophoblast stem cells"

**Supplemental Tables**

Table S1. FPKM, DEG (differentially expressed gene) analysis, z-score cluster analysis of hypoxia RNA-seq samples. Related to Figure 1.

Table S2. List of 100 specific cell type genes corresponding to FPKM of hypoxia samples. Related to Figure 2.

Table S3. FPKM for Control and GCM1-/- samples and DE-seq analysis. Related to Figure 2.

Table S4. Coordinates of ChIP-seq and ATAC-seq peak sets. Related to Figure 3.
